# Ensemble-specific deficit in neuronal intrinsic excitability in aged mice

**DOI:** 10.1101/2022.06.02.494525

**Authors:** Lingxuan Chen, Taylor R. Francisco, Austin M. Baggetta, Yosif Zaki, Steve Ramirez, Roger L. Clem, Tristan Shuman, Denise J. Cai

## Abstract

With the prevalence of age-related cognitive deficits on the rise, it is essential to identify cellular and circuit alterations that contribute to age-related memory impairment. Increased intrinsic neuronal excitability after learning is important for memory consolidation, and changes to this process could underlie memory impairment in old age. Some studies find age-related deficits in hippocampal neuronal excitability that correlate with memory impairment but others do not, possibly due to selective changes only in activated neural ensembles. Thus, we tagged CA1 neurons activated during learning and recorded their intrinsic excitability 5 hours or 7 days post-training. Adult mice exhibited increased neuronal excitability 5 hours after learning, specifically in ensemble (learning-activated) CA1 neurons. As expected, ensemble excitability returned to baseline 7 days post-training. In aged mice, there was no ensemble-specific excitability increase after learning, which was associated with impaired hippocampal memory performance. These results suggest that CA1 may be susceptible to age-related impairments in post-learning ensemble excitability and underscore the need to selectively measure ensemble-specific changes in the brain.

## 1 Introduction

Aging is associated with functional decline in some aspects of cognitive performance and brain function, including memory difficulty in older adults (Luo and Craik, 2008; Newson and Kemps, 2006). In particular, hippocampus-dependent memories show age-related impairment both in humans (Newman and Kaszniak, 2000) and rodents (Barnes, 1979; Gallagher et al., 2015; Kwapis et al., 2019; Rapp et al., 1987; Wimmer et al., 2012). To move towards ameliorating age-related memory impairment, some studies have focused on hippocampal neurophysiological changes during aging. One neuronal property that supports memory processing is intrinsic excitability—a neuron’s tendency to fire action potentials upon synaptic integration, primarily dictated by the expression and function of voltage-gated ion channels on the cell membrane. Intrinsic excitability is crucial for memory processes such as memory allocation, consolidation, and updating (Chen et al., 2020; Daoudal and Debanne, 2003; Mau et al., 2020; Sweis et al., 2021; Zhang and Linden, 2003). After learning, neuronal intrinsic excitability of hippocampal CA1 pyramidal cells temporarily increases and returns to baseline within 7 days (Moyer et al., 1996). Post-learning increases in CA1 neuronal intrinsic excitability are observed across memory paradigms (Kaczorowski and Disterhoft, 2009; Matthews et al., 2009; Mckay et al., 2009; Oh et al., 2003; Song et al., 2012; Zelcer et al., 2006); particularly, it may support the stabilization of newly acquired information after learning, i.e. memory consolidation (Ryan et al., 2015) (see Discussion).

Young animals that successfully learn exhibit increased excitability of CA1 pyramidal cells, which is not observed in aged animals that fail to learn (Disterhoft and Oh, 2007; Mcechron et al., 2001; Moyer et al., 2000; Tombaugh et al., 2005). However, other studies found no change in CA1 during normal aging (Small et al., 2004; Wilson, 2005; Yassa et al., 2011). Since only a sparse cell population, a neural “ensemble”, is involved in memory encoding (Guzowski et al., 1999; Lacagnina et al., 2019; Reijmers et al., 2007), excitability changes underlying memory encoding may be specific to learning-activated ensemble neurons. Indeed, activity-dependent labeling techniques (Denny et al., 2014; Guenthner et al., 2013; Reijmers et al., 2007) demonstrate that ensemble cells have increased excitability compared to neighboring non-ensemble cells after learning (Cai et al., 2016; Crestani et al., 2018). Furthermore, ensemble cells have been shown to be crucial for memory consolidation (Ryan et al., 2015) and disrupting ensemble-specific reactivation shortly after learning impairs consolidation (Hsiang et al., 2014; Park et al., 2016). In aged rats, the expression level of immediate early gene *Arc* was reduced in CA1 after exploration (Penner et al., 2011), suggesting lower post-learning neuronal activity relative to younger brains. Yet, the extent to which normal aging alters post-learning excitability and whether it does so in a general or ensemble-specific manner remains unclear. Here, we find that aging specifically affects post-learning excitability in hippocampal CA1 ensemble cells, likely leading to memory impairment via disrupted memory consolidation. Interestingly, neuronal excitability of CA1 non-ensemble cells did not change with aging, suggesting the specificity of this deficit. Our work resolves prior discrepancies by using activity-dependent tagging to reveal that ensemble-specific excitability increase after learning is absent in memory-impaired older mice. Our study demonstrates that CA1 ensemble cells may be especially susceptible to age-related excitability deficits and underscores the importance of using ensemble-specific techniques to investigate changes underlying age-related memory impairment.

## 2 Materials and Methods

### 2.1 Animals

All experiments were approved in advance by the Institutional Care and Use Committee of the Icahn School of Medicine at Mount Sinai and conducted on young (3-6 months) and old (22-24 months) C57Bl/6J male mice obtained from the Jackson Laboratory (Bar Harbor, ME). Mice were housed 4-5/cage with ad libitum food and water on a 12-hour light-dark cycle (lights on at 0700 hours).

### 2.2 Novel Object Location Task

Prior to all experiments, mice were handled in the vivarium for 1 minute per day for 4 days and habituated to transportation and external environmental cues for another 4 days. Mice were also habituated to empty arenas (1 ft X 1 ft) with spatial cues on the walls for 10 minutes over 4 days. On the training day, mice explored in arenas with 2 identical objects (2-inch-tall plastic trophies) taped to the upper corners of the arenas to prevent changing of objects’ locations during animal exploration and interaction. Young adult mice spent 10 minutes in the arenas. To control for interaction differences, each aged mouse was randomly paired with a young adult mouse and explored the arena until the total interaction time with both objects reached that of the paired young mouse. One day later (test day), mice spent 5 minutes in the same arena with the same objects but with 1 object moved to a new location (Fig. 1A). Exploration was digitally recorded and hand scored for time spent exploring objects. Interaction was defined as active sniffing or physical contact with the objects. Mice that spent less than 30 seconds interacting with objects or less than 8 minutes in the arena during the training session were excluded. Discrimination index (DI) was calculated as:

**Figure 1.**
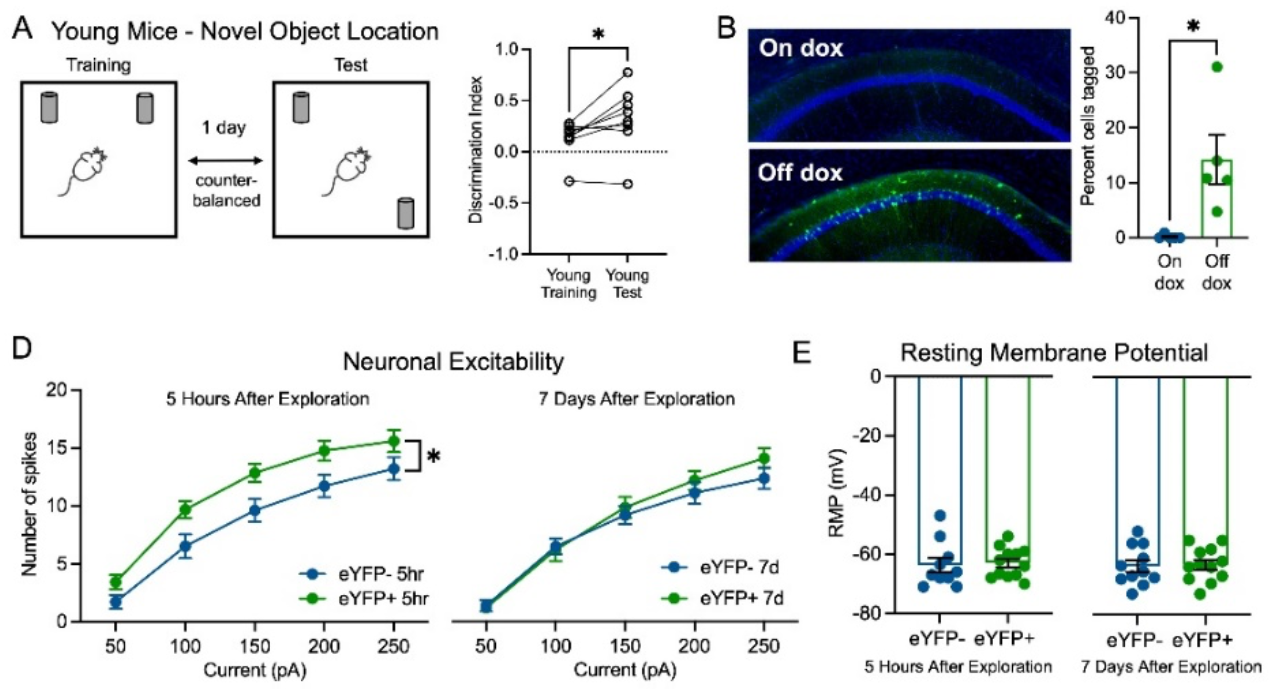
Young adult mice with intact memory in the novel object location task had transiently increased neuronal excitability in CA1 ensemble cells after learning. (A) Schematic of novel object location task. Young mice had higher DI in test session vs. training session (paired t-test, p=0.03, N=8). (B) Doxycycline (dox) controls the tagging of ensemble cells in CA1. Left: representative images of eYFP+ in CA1 in mice on and off dox. Right: percentage of cells tagged in both groups (unpaired t-test, p=0.01, N=5 mice/condition). (C) In young adult mice (3-6 months), neuronal excitability of CA1 ensemble cells increased 5 hours after context exploration compared to non-ensemble cells (two-way RM ANOVA, F_1,20_=5.68, p=0.03, n=10 cells eYFP-, 12 cells eYFP+). Neuronal excitability of ensemble cells returned to the same level as non-ensemble cells 7 days after (two-way RM ANOVA, F_1,21_=0.32, p>0.05, n=11 cells eYFP-, 12 cells eYFP+). (D) In young mice, the resting membrane potential (RMP) of ensemble cells was not different from that of non-ensemble cells 5 hours (unpaired t-test, p>0.05, n=10 cells eYFP-, 12 cells eYFP+) or 7 days (unpaired t-test, p>0.05, n=11 cells eYFP-, 12 cells eYFP+) after learning.

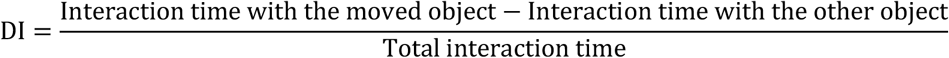

### 2.3 Virus infusion

We used an activity-dependent virus cocktail of AAV9-cFos-tTA (titer: 3×10^13^ GC ml^-1^) and AAV9-TRE-eYFP (titer: 1×10^13^ GC ml^-1^) that labels neurons expressing the immediate early gene *cFos* in a doxycycline (dox)-dependent manner (Chen et al., 2019; Cincotta et al., 2021). Mice were anaesthetized with 1.5 to 2.0% isoflurane for surgical procedures and placed into a stereotactic frame (David Kopf Instruments, CA). Lidocaine (2%) was applied to the sterilized incision site as an analgesic; carprofen (5 mg kg−1) and ampicillin (20 mg kg^-1^) were administered immediately after surgery and for 6 subsequent days. Mice were bilaterally injected with 300 nL of virus cocktail at 2 nl sec^-1^ into dorsal CA1 using coordinates: −2.0 mm AP, ±1.5 mm ML, −1.5 mm DV, all from bregma, using a Nanoject III injector (Drummond Scientific Company, PA). Mice were put on dox chow (1 g kg^-1^, Bio-Serve, MD) immediately after surgery to inhibit cell labelling.

To validate dox-dependent eYFP expression (Fig. 1B), young mice (4 months) were handled and habituated to the behavior room over 10 days after virus infusion. To activate labelling, a subset of mice was taken off dox and given regular chow for 48 hours (“off dox”), while others remained on 1 g kg^-1^ dox (“on dox”). Mice were introduced to a novel context (Med Associates Inc., VT) and allowed to freely explore for 10 minutes. Mice were immediately given 1 g kg^-1^ dox chow to prevent further cell labelling and sacrificed 24 hours later. Brains were dissected 24 hours after the context exploration or homecage and post-fixed in 4% paraformaldehyde at 4°C for 24 hours, coronally sliced at 70 μm using a vibratome (Leica VT1000S, Leica Biosystems, IL), and mounted with DAPI Fluoromount-G (SouthernBiotech, AL). Images were taken with an epifluorescence microscope (Leica DM6 B, Leica Biosystems, IL). An experimenter blind to the groups manually counted the numbers of eYFP+ and DAPI cells in one region from one slice per mouse. The selected region was a 1 mm x 0.5 mm rectangle box (in mm: ML +/-1.5±0.5, DV −1.25±0.25, at AP −2.0, all relative to Bregma). Only cells in the CA1 pyramidal cell layer were counted.

### 2.4 Whole-cell patch clamp recordings in acute brain slices

Behaviorally naïve mice were introduced to a novel context (Med Associates Inc., VT) and allowed to freely explore for 10 minutes. Two littermates explored the novel contexts on the same day, then randomly assigned to the 5-hour and 7-day groups for patch clamp experiments. Five hours or seven days later, mice were anesthetized by isoflurane through inhalation, followed by rapid decapitation. Coronal acute brain slices were prepared on a vibratome (Leica VT1200S, Leica Biosystems, IL) at 350 μm thickness in ice-cold cutting solution. Slices were kept in a submerged chamber filled with sucrose-based artificial CSF (sACSF). The recovery chamber was kept in 37°C water bath for 30 min and then moved to room temperature. For whole-cell recording, acute brain slices were placed in a submerged chamber filled with circulating ACSF at room temperature. A K-met based intracellular solution was used for recording excitability measurements. The recipes of all solutions can be found in a previous publication from the lab (Shuman et al., 2020). Cells were visualized on a Nikon Eclipse FN1 microscope (Nikon Instrument Inc., NY) paired with a SOLA light engine (Lumencor, OR) to identify non-fluorescent and eYFP-labeled cells. Principal excitatory neurons were selected based on pyramidal morphology under DIC microscopy and physiology. Recordings were conducted under current clamp mode using a Multiclamp 700B amplifier and an Axon Digidata 1550B digitizer (Molecular Devices Inc., CA). Resting membrane potential was measured 3 minutes after transitioning into the current-clamp mode. Spike-current curves were determined by injecting a series of current steps (0-300 pA at 50 pA increments, 500 ms). Data analysis was done using Clampfit 10.7 (Molecular Devices Inc., CA). The numbers of non-ensemble and ensemble cells recorded in each young and aged mouse at each time point are shown in Supplementary Table 1.

### 2.5 In vivo calcium imaging with Miniscopes

Using the TetTag system, dox was taken off for 48 hours, resulting in tagging cells that are active in the homecage when there was low or no dox in the system. In order to estimate the fraction of spontaneously active cells while the animals are in their homecage that may become active during subsequent learning, we performed *in vivo* calcium imaging with Miniscopes in a separate cohort of 4 young adult mice (4-5 months). Although there are differences between cFos and calcium, cells that have high calcium activity are more likely to upregulate cFos protein than cells that have low calcium activity (Pettit et al., 2022). We imaged calcium activity for 10 min while the animals were in their homecage and during their subsequent context learning 36 hours later. We quantified the proportion of cells that were reactivated during contextual learning among all cells active during the 10 min homecage session (Supplement Fig. 1).

#### Surgery

Mice underwent three serial surgeries: 1) to express the calcium indicator GCaMP6f, 2) to implant a GRIN lens above hippocampus, and 3) to attach a baseplate for Miniscope recording. During the first surgery, 300 nl of AAV1-Syn-GCaMP6f-WPRE-SV40 (at ∼2×10^12^ GC ml^-1^) was injected into dorsal CA1 (AP: −2 mm, ML: −1.5 mm, DV: −1.2 mm relative to bregma) at 2 nl sec^-1^ and allowed 5min for diffusion time, after which the incision was sutured. Two weeks later, mice had their overlying cortex aspirated and a 1mm x 4mm (diameter x length) GRIN lens implanted 200um above the injection site, using dental cement. Two weeks later, mice underwent a third surgery where a baseplate was implanted visually guided by a Miniscope to find the field of view which optimally resolved neuronal cell bodies.

#### Miniscope Habituation/Recordings

Mice were housed in a custom-made homecage (Maze Engineers) with a food hopper and water bottle mounted on a side wall, and with a slit along the ceiling to accommodate a Miniscope coaxial cable. Mice had the Miniscope chronically attached for the duration of the experiment and were allowed about one week to adjust to the weight of the Miniscope as well as living in the new homecage. The Miniscope was connected to a coaxial cable (Cooner Wire) which connected to a low-torque commutator (Neurotek) to allow the mice to freely move around the homecage with minimal rotational force. 36 hours prior to context exposure, mice underwent a 10 min calcium imaging recording to sample their spontaneous neuronal activity in their homecage. 36 hours later, mice were exposed to a novel context with unique visual, tactile, and olfactory cues (Med Associates), during which calcium activity was also recorded.

#### Data Processing/Analysis

To extract activity of individual neurons, we used Minian, an open-source tool which corrects for background fluorescence and motion before using a constrained non-negative matrix factorization approach to extract calcium transients of individual cells (Dong et al., 2022). For each mouse, we cross-registered cells across recordings using CellReg, an open-source tool which registers cells across sessions based on their location and spatial correlation with one another, after aligning recordings based upon the spatial footprints of the tracked cells (Sheintuch et al., 2017). To analyze the resulting data, we used custom-written Python scripts which we will share upon request.

## 3 Results

### 3.1 Young mice with intact hippocampal memory had transient increase in excitability in CA1 ensemble cells

We used a novel object location task to test hippocampal-dependent memory (Murai et al., 2007; Wimmer et al., 2012). Young adult mice initially interacted similarly with the two objects (Fig. 1A; average DI=0.13, one sample t-test, p>0.05). When returned to the arena 1 day later with 1 object moved (Fig. 1A), young mice showed a preference for the newly located object (average DI=0.33, one sample t-test, p=0.02), indicating a memory for the old location and suggesting intact hippocampal-dependent memory.

To determine the time course of excitability changes in CA1 ensemble cells, we first confirmed that doxycycline (dox) effectively controlled the expression of eYFP in dorsal CA1 (Fig. 1B). Five hours after learning, tagged CA1 ensemble neurons (eYFP+) displayed significantly higher intrinsic excitability relative to non-ensemble cells, firing more action potentials when injected with the same amount of current (Fig. 1C). Excitability of ensemble cells returned to non-ensemble levels within 7 days (Fig. 1C), demonstrating the transiency of post-learning excitability increase. Ensemble-specific plasticity was not due to changes in resting membrane potential (RMP), as young mice exhibited no difference in RMP of ensemble and non-ensemble neurons either 5 hours or 7 days after learning(Fig. 1D), consistent with previous work (Crestani et al., 2018; Mckay et al., 2009; Moyer et al., 2000, 1996; Song et al., 2012). These results demonstrate that young adult mice with intact hippocampus-dependent memory exhibit transient, ensemble-specific excitability increase in CA1 after learning.*3*.*2 Impaired hippocampal memory and lack of increased excitability in CA1 ensemble cells of aged mice*

As with young adult mice, aged mice interacted equally with both objects during training (Fig. 2A, average DI=0.07, one sample t-test, p>0.05). During test, there was no difference in the total interaction time with the two objects between aged and young mice (Fig. 2B, unpaired t-test, p>0.05), however aged mice did not exhibit a preference for the moved object (Fig. 2A, average DI=0.19, one sample t-test, p>0.05), indicating impaired hippocampus-dependent memory in aged mice, consistent with previous work (Murai et al., 2007; Wimmer et al., 2012). In contrast to young counterparts, aged mice did not show increased intrinsic excitability of ensemble neurons 5 hours after learning (Fig. 2C). As with young mice, we found no differences in RMP between ensemble and non-ensemble neurons in aged mice (Fig. 2D). These results indicate that aging specifically affects the post-learning increase in the neuronal intrinsic excitability of ensemble cells.

**Figure 2.**
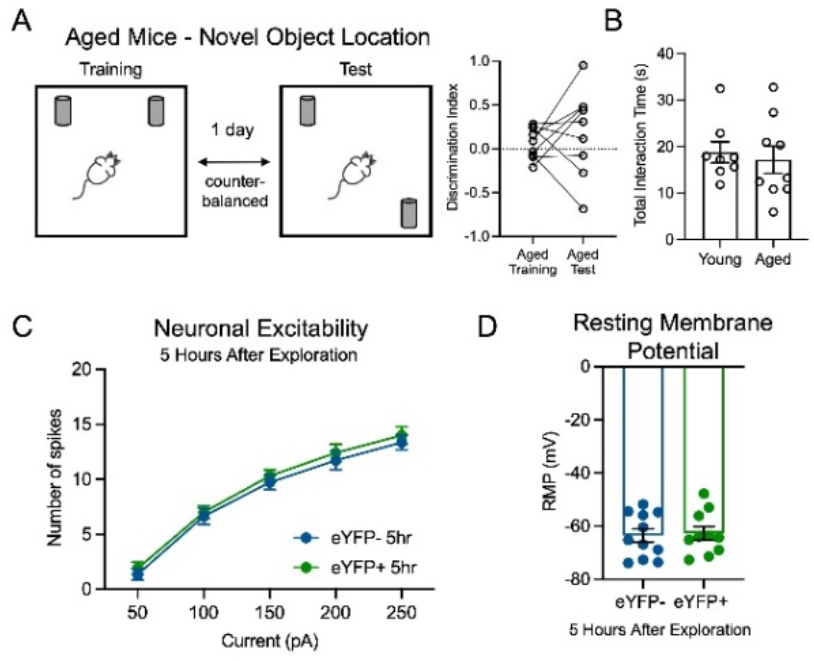
Aged mice with impaired memory in a novel object location task had no increase in neuronal excitability of CA1 ensemble cells after learning. (A) Aged mice had comparable DI in novel object test and training sessions (paired t-test, p>0.05, N=9). (B) Total interaction time with the two objects did not significantly differ between aged and young mice (N=8 and 9, young and aged, respectively, unpaired t-test, p>0.05). (C) In aged mice, neuronal excitability of CA1 ensemble cells stayed comparable to non-ensemble 5 hours after context exploration (two-way RM ANOVA, F1,19=0.40, p>0.05, n=11 eYFP-, 10 eYFP+). (D) In aged mice, the RMP of ensemble cells was not different from non-ensemble cells 5 hours after learning (unpaired t-test, p>0.05, n=11 eYFP-, 10 eYFP+).

## 4 Discussion

We find that normal aging particularly affects neuronal intrinsic excitability in ensemble cells shortly after learning. As predicted, younger mice exhibited temporarily increased excitability of ensemble cells hours after learning, which returned to baseline days later. Aged mice with impaired hippocampal memory performance exhibited deficit in post-learning excitability of ensemble cells. These results indicate that with normal aging, CA1 is susceptible to impairment in the regulation of post-learning excitability in ensemble cells, which may lead to disrupted memory consolidation and hippocampus-dependent memory deficits.

Neuronal intrinsic excitability is measured primarily using *in vitro* whole-cell patch clamp recordings in brain slices (Disterhoft and Oh, 2007). Studies have also used *in vivo* firing frequency (Mcechron et al., 2001; Wilson, 2005) or BOLD brain activity (Small et al., 2004; Yassa et al., 2011) as a proxy for neuronal excitability. Conflicting results related to intrinsic excitability in CA1 pyramidal cells during aging may arise from a) differing methods and parameters, b) differing behavior paradigms and excitability measurement time points, and/or c) the population of cells recorded. We have shown that age-related post-learning excitability changes are specifically isolated to ensemble cells, resolving the discrepancy in previous studies. An ensemble-specific approach is more sensitive to age-related cellular and circuit alterations contributing to memory processing. We note, however, that cFos-driven tagging (Chen et al., 2019; Cincotta et al., 2021; Reijmers et al., 2007) does not capture all cells activated during learning—active cells may not have activated cFos or may express other immediate early genes (Sun et al., 2020), and that cells not tagged by this system may also be involved in memory formation.

The transient increase in neuronal intrinsic excitability is crucial for memory consolidation. Intrinsic excitability and long-term potentiation, a well-established proxy of memory consolidation (Kandel et al., 2014), share common voltage channels and signaling pathways (Chen et al., 2020), indicating that the processes are closely correlated and one may enhance the other. Lack of learning-induced excitability increase in certain brain regions correlated with behavior deficits, and pharmacologically increasing excitability in these regions rescued behavior (Cai et al., 2016; Disterhoft and Oh, 2006; Yu et al., 2017). Therefore, enhanced intrinsic excitability in hippocampal ensemble cells may initially support synaptic consolidation in the local circuit hours after learning, and increased excitability lasting days may further support cross-regional synaptic connections, thus contributing to systems consolidation on a longer time scale. In normal aging, post-learning excitability deficits in ensemble cells may disrupt stabilization of newly acquired information, leading to impaired memory. Given the importance of ensemble-specific excitability increase during memory consolidation, it will be important for future studies to monitor neuronal excitability of ensemble cells during the consolidation phase using *in vivo* techniques with high temporal and spatial resolution, such as fluorescent voltage sensors (Piatkevich et al., 2019).

## Acknowledgement

This work was supported by the Brain Research Foundation Award, Klingenstein-Simons Fellowship, NARSAD Young Investigator Award, McKnight Memory and Cognitive Disorder Award, One Mind-Otsuka Rising Star Research Award, Mount Sinai Distinguished Scholar Award, DP2 MH122399-01, and R01 MH120162-01A1 to DJC; the CURE Taking Flight Award, American Epilepsy Society Junior Investigator Award, R03 NS111493, R21 DA049568, R01 NS116357, and RF1 AG072497 to TS; R01 124880 and R01 116445 to RLC; and an NIH Early Independence Award DP5 OD023106-01, NIH Transformative R01 Award, a Ludwig Family Foundation grant, and McKnight Foundation Memory and Cognitive Disorders Award to SR. We thank Stellate Communications for assistance with the preparation of this manuscript.

