## Supplementary Figure 1 for "Ensemble-specific deficit in neuronal intrinsic excitability in aged mice"

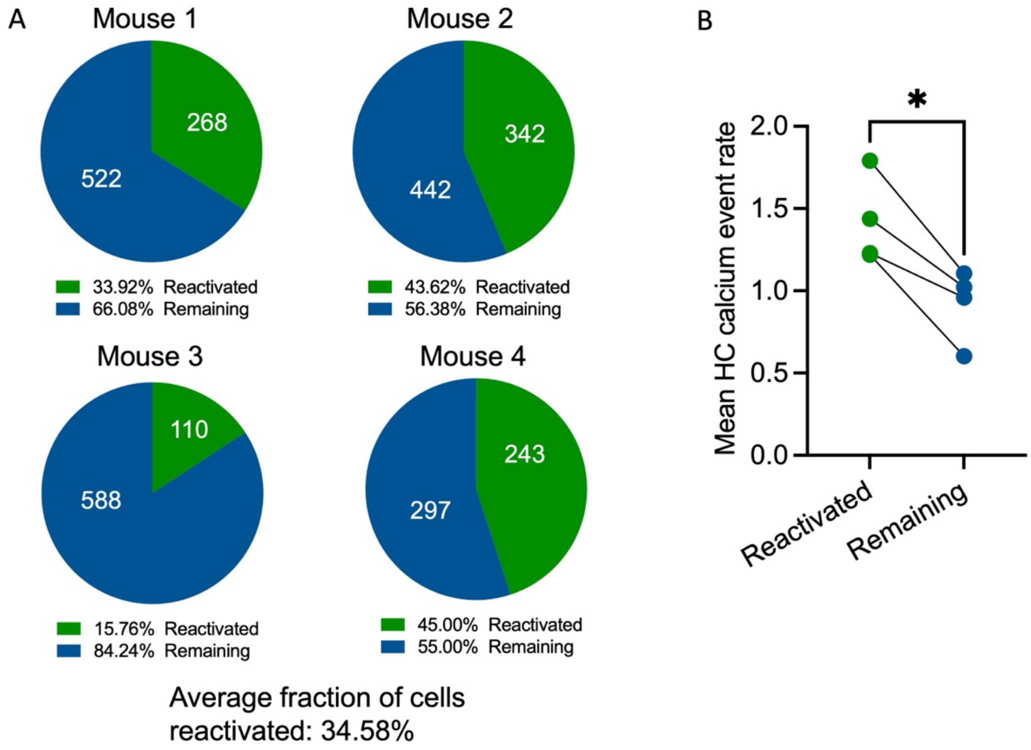


**Supplementary Figure 1.** *In vivo* calcium imaging showing that highly active cells in homecage were reactivated during contextual learning. (**A**) In each mouse (total N=4), the fraction of active cells in homecage (over 10 min, 36 hours prior to contextual learning) that were reactivated during the 10 min learning session (green: reactivated; blue: remaining; white: number of cells). An average of 34.58% of cells active in the homecage were reactivated during contextual learning 36 hours later. (**B**) Reactivated cells were more active in homecage compared to the remaining cells (N=4 mice, paired t test, reactivated vs. remaining, p=0.01), consistent with previous literature showing that cells that are more excitable prior to a novel experience are more likely to be recruited into the memory ensemble (Cai et al., 2016; Han et al., 2007; Park et al., 2016; Yiu et al., 2014; Zhou et al., 2009). Given that cells that fire more, or have more calcium activity, are more likely to express cFos (Pettit et al., 2022; Tanaka et al., 2018), the TetTag system is likely tagging many of these highly active cells in the homecage, and these active cells are also more likely to be reactivated during subsequent learning.
