## Supplementary Table 1 for "Ensemble-specific deficit in neuronal intrinsic excitability in aged mice"

**Supplementary Table 1**. Numbers of cells recorded in each young and aged mouse at each time point.

| Young 5 hr | | | Young 7 d | | | Aged 5 hr | | |
|---|---|---|---|---|---|---|---|---|
| Mouse ID | eYFP- cells | eYFP+ cells | Mouse ID | eYFP- cells | eYFP+ cells | Mouse ID | eYFP- cells | eYFP+ cells |
| Y_01 | 2 | 3 | Y_08 | 2 | 1 | A_01 | 1 | 2 |
| Y_02 | 1 | 0 | Y_09 | 1 | 2 | A_02 | 1 | 1 |
| Y_03 | 3 | 0 | Y_10 | 0 | 1 | A_03 | 3 | 2 |
| Y_04 | 0 | 1 | Y_11 | 2 | 4 | A_04 | 1 | 1 |
| Y_05 | 2 | 3 | Y_12 | 3 | 4 | A_05 | 1 | 2 |
| Y_06 | 0 | 2 | Y_13 | 3 | 0 | A_06 | 0 | 2 |
| Y_07 | 2 | 3 | | | | A_07 | 4 | 0 |
| **Total** | 10 | 12 | **Total** | 11 | 12 | **Total** | 11 | 10 |
